# Intranasal AAV Vaccination of SARS-CoV-2 Induce Strong and Sustained Neutralizing Antibodies in Mice

**DOI:** 10.1101/2025.03.14.640671

**Authors:** Youngmi Ji, Giovanni Di Pasquale, Changyu Zheng, Sandra Afione, Thomas Esperanza, Hongen Yin, Peter D. Burbelo, John A. Chiorini

**Affiliations:** Adeno-Associated Virus Biology Section, National Institute of Dental and Craniofacial Research, National Institutes of Health, Bethesda, MD 20892; National Institute of Neurological Disorders and Stroke, Bethesda, MD, USA

**Author notes:** Both contributed equally (Youngmi Ji, Giovanni Di Pasquale).

## Abstract

The COVID-19 pandemic continues to pose significant health challenges, despite existing vaccines. This study evaluates the immunogenicity of recombinant adeno-associated viruses (AAV) expressing SARS-CoV-2 spike proteins, administered intramuscularly and intranasally in mice. Both delivery methods of AAV5-spike, AAV5-spike stabilized trimer as well as AAV44.9-spike elicited robust serum anti-spike antibodies within 8-12 weeks, with high levels of anti-spike antibodies sustained for over a year. Comparison of mouse serum antibodies 16 weeks post intramuscular or intranasal AAV5 administration demonstrated similar SARS-CoV-2 spike binding neutralizing activity in vitro. Analysis of changes in cellular immunity by ELISpot at 12 weeks post-AAV spike transduction revealed interferon-γ induction in response to peptide challenge. Despite a decline in AAV vector DNA at the injection site, the persistence of anti-spike antibodies demonstrated that AAV-vectors can elicit lasting immune responses, highlighting nasal AAV administration as a potential strategy to block respiratory virus infections.

## Introduction

Coronavirus disease 2019 (COVID-19) caused by the SARS-CoV-2 virus infection has resulted in a devastating toll marked by a high rate of respiratory-related mortality and additional morbidity involving injury to the lungs and kidney, cognitive dysfunction, and chronic fatigue. The variability in clinical symptoms with COVID-19 likely involves different sites of SARS-CoV-2 localization, and host factors including genetic differences, age, gender, and coexistence of other comorbid conditions. In late 2020, adenovirus-based and mRNA-based COVID-19 spike protein vaccines represented a significant breakthrough, substantially reducing mortality and hospitalization following SARS-CoV-2 infection [1]. These new vaccine platforms facilitated rapid and cost-effective manufacturing of COVID-19 vaccines and were instrumental in blunting the pandemic. Despite the success of the COVID-19 vaccines in blocking systemic infection, they do not provide robust and sustained mucosal immunity [2, 3]. The limited mucosal immunity elicited by current COVID-19 vaccines explains the lack of efficacy in blocking SARS-CoV-2 infection and onward viral transmission and spread. SARS-CoV-2 may replicate in the airway for some time before effective IgG-mediated neutralization occurs, allowing transmission during that period [4]. The induction of long-lasting mucosal immunity required to block initial infection remains an active area of interest. Moreover, the continued evolution of SARS-CoV-2 necessitates the adaption of these new variants into vaccines which presents additional challenges [1].

Adeno-associated vectors (AAVs) for gene therapy are highly successful in treating several monogenic diseases and have a demonstratable large safety record [5]. Besides the treatment of genetic diseases, AAV gene delivery technologies have also been used to express therapeutic monoclonal antibodies against infectious agents [6] and for the delivery of experimental vaccines against such pathogens as HIV [7] and influenza [8, 9]. More recently, several reports have also used the AAV platform to induce SARS-CoV-2 spike vaccine immunizations in both mice [10–14] and macaques [10–12, 15]. Many of these studies utilized intramuscular (IM) delivery of the SARS-CoV-2 spike vaccine formulation. For example, Zabaleta et al. showed in both mice and macaques that IM administration of an AAV-Spike can induce potent neutralizing antibodies and cell-mediated immunity against SARS-CoV-2 [10]. Based on numerous successful AAV studies showing intranasal (IN) administration of transgene expression [16], as well as, our previous AAV12 study documenting the IN delivery of influenza antigen for specific vaccine immune responses [17], we hypothesized that an IN AAV-gene delivery strategy might be particularly suited to inducing durable SARS-CoV-2 spike mucosal immunity.

In the current study, the ability of three different AAVs (AAV5, AAV-44.9, AAV-12) expressing SARS-CoV-2 spike and/or spike trimer antigen configurations administered by the IM and/or IN routes were evaluated for *in vivo* transduction of mice for mounting immune responses against the spike protein. The different AAV vectors encoding SARS-CoV-2 spike proteins delivered by a single IM and IN administration resulted in high levels of sustained antibody production for over a year and additional cellular immunity. Our findings suggest that numerous AAVs show efficacy in eliciting effective humoral vaccination against SARS-CoV-2 and the intranasal administration may be a useful immunization route for vaccination against SARS-CoV-2 and other respiratory viruses.

## Results

### AAVs Transgenes Design for SARS-CoV-2 Spike and Routes of Mice Administration

Three different AAV types, AAV5, AAV44.9, and AAV12, were tested for SARS-CoV-2 spike protein vaccine immunization (Table I). These isolates were chosen based on their diverse attachment receptors and distinct transduction activity. Two different SARS-CoV-2 spike protein configurations, the entire ectodomain of the spike protein from amino acids 1-1208 and a Spike-trimer stabilized form were generated for AAV expression. To generate these recombinant constructs, two AAV ITR expressing cassettes containing either the SARS-CoV2 spike (the entire ectodomain from amino acids 1-1208) and a spike-trimer stabilized form, both driven by a CMV promoter were built. The design of each of the cassettes is shown in Supplemental Figure 1A. Transfection of the CMV expression plasmids encoding both the spike and spike-trimer into HEK293 along with Western blotting of cell lysate and supernatant detected recombinant spike protein of 160 KDa (Supplemental Figure 1B). Following AAV packaging and purification, 5 × 10^11^ DNase-resistant particles AAVs were administered to mice by two different routes, IM and IN in a volume of 50 or 150 μl of PBS respectively.

**Table I.** Characteristics of the AAV serotypes, SARS-CoV-2 spike proteins and Routes of AAV.

| AAV serotype | SARS-CoV-2 Design | Administration |
| --- | --- | --- |
| AAV5 | Spike | IM |
| AAV5 | Spike-Trimer | IM |
| AAV5 | Spike | IN |
| AAV5 | Spike-Trimer | IN |
| AAV44.9 | Spike | IN |
| AAV12 | Spike | IN |

### Humoral Responses to AAV-Mediated SARS-CoV2-Spike Transduction

Female mice were vaccinated with the AAV-spike and AAV-spike trimer by IM and IN routes. During the course of the study for over a year, all the mice for monitoring antibody levels following AAV transduction survived, including control untransduced mice. The levels of anti-spike IgG were detected by the LIPS assay and quantitatively expressed as relative light units (LU) values. One week after IM administration, the presence of low levels of seropositive spike antibodies were already detected in the range of 20,000-80,000 LU in the sera of several mice treated with AAV5-spike and AAV5-spike trimer (Figure 1A). By 12 weeks, high levels of spike antibody (>1,000,000 LU), similar to the antibody levels seen in natural human infection or human vaccination were detected in the sera of all mice who received either AAV5-spike or AAV5-spike trimer (Figure 1A). Longitudinal anti-spike antibody responses were also observed in the mice who received AAV5-spike, AAV5-spike trimer, and AAV 449-spike by IN administration (Figure 1B). No statistical difference was seen when comparing the antibody levels observed over time by AAV5-spike comparing IM and IN administration. An additional group of mice used AAV12-spike administrated intranasally also showed a similar antibody profile against the spike protein (Supplemental Figure 2). Remarkably, over the course of a year after AAVs IM or IN administration, the mice continued to show high levels of anti-spike antibodies with little diminution over time (Figure 1A and 1B and Supplemental Figure 2). Although no difference in antibody levels was noted between using either AAV5-spike protein compared to the AAV5-spike protein trimer administered IN (Figure 1B), the AAV5-spike trimer tracked towards lower levels than AAV5-spike by IM administration (Figure 1A). Taken together the data shows that spike antibodies levels generated by AAVs treatment were as high as levels produced during natural SARS-CoV-2 infection and after human vaccination and remained high for over one year.

**Figure 1.**
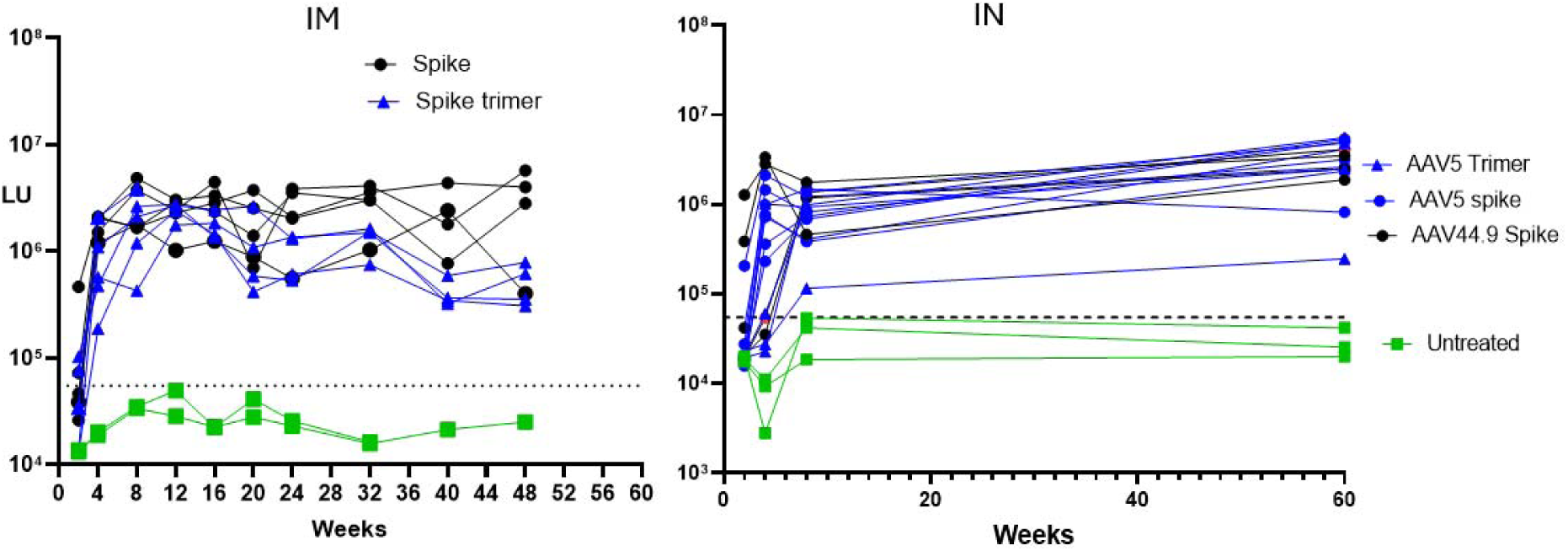
SARS-CoV-2 Spike Antibody Levels by LIPS in Treated Mice at Different Times post IM and IN After AAVs Administration. Each line represents a value from each serum from mice over time. As control, untreated serum mice were also quantified and shown individually. Antibody levels are expressed in relative light units (LU) on a log scale. The dashed line represents the cut-off value for determining spike antibody seropositivity.

### Serum from Treated Mice Results in SARS-CoV-2 Spike Neutralizing Activity

To assess the antiviral activity of SARS-CoV-2 antibodies secreted in mice, the *in vitro* competition assay was performed to determine the level of neutralizing antibodies. This *in vitro* assay mimics the first step of the interaction of the SARS-CoV-2 natural infection with the spike viral protein interacting with the target cellular Ace2 receptor. Based on the high levels of antibody detected at 16 weeks by LIPS, serum samples from this cross-sectional time point serum samples were tested for neutralizing antibodies from the 3 groups of mice treated with AAV-spike along with untreated control mice. As expected, the control mice showed no significant neutralizing activity (Figure 2), whereas all but one of the mice from IM treated with AAV5-spike and spike trimer showed close to maximum neutralizing activity against the SARS-CoV-2 respectively (Figure 2). Interestingly, all the mice receiving IN administration of AAV5-Spike and AAV5-spike trimer showed maximum neutralizing activity (Figure 2). Nevertheless, no statistical neutralizing activity difference could be detected between the IM and IN-treated mice, and between mice administered AAV-spike or AAV5-spike trimer. Overall, these results highlight the potent neutralizing activity of serum at 16 weeks post-treatment either with AAV5-spike or AAV5-spike trimer by both routes of administration.

**Figure 2.**
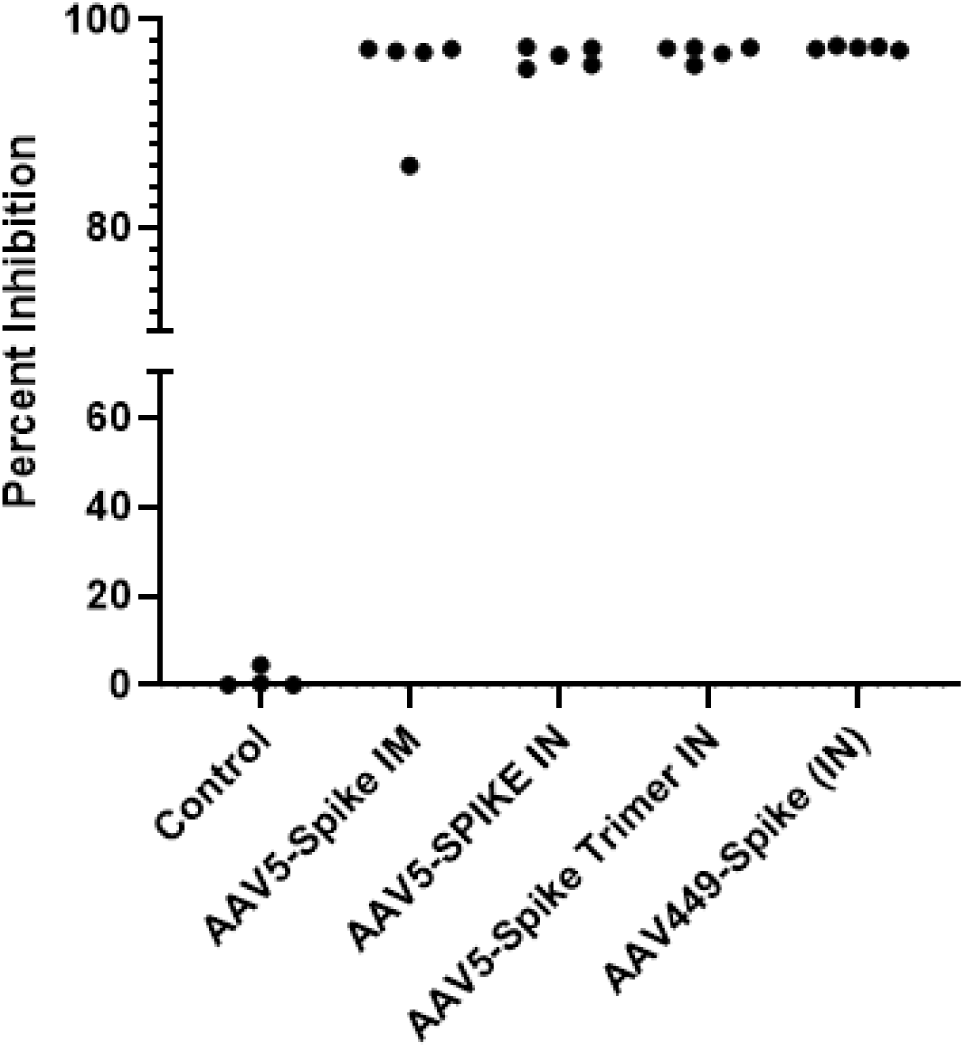
Neutralizing Activity of Mouse Serum Anti-Spike Antibody After 16 Weeks Post-Administration. Mouse sera collected from control or mice transduced with AAV-spike by the IM or IN route were tested for *in vitro* neutralizing activity in blocking interactions between spike-Ace proteins as described in the material and methods. Each data point represents an individual mouse serum from each of the five groups of mice tested. Statistical analysis by Mann Whitney *U* test revealed highly statistical (P ≤0.001) neutralizing activity in the mice receiving AAV-spike by IM and IN administration.

### Cellular Immunity Against Spike in AAV Transduced Mice

Based on strong humoral responses observed against SARS-CoV-2 spike protein of AAV-spike treated mice, the immunological analysis was extended to study cellular immunity. For these experiments, a separate group of mice were transduced with either AAV5-GFP or AAV5-spike for studying T-cells responses. For these experiments, two mice were given AAV5-GFP, and four mice received AAV5-spike by IM administration and 16 weeks later, the mice were sacrificed and splenocytes were isolated from the corresponding mice. Using ELISpot, the splenocytes from the mice were challenged with a pool of SARS-CoV-2 spike peptides. The AAV5-GFP control mice by ELISpot assay showed a very low induction of IFN-γ production (4 spots/10^6^ PBMC) (Figure 3). In contrast, AAV5-spike-treated mice generated significantly higher levels of IFN-γ production (Figure 3). However, there was variability among treated mice, in which one mouse demonstrated high levels of IFN-γ (232 spots/10^6^ PBMC) resulting in about 1/15 of the potent signal seen with PMA stimulation. The three other mice had values less pronounced production generating approximately 30 spots/10^6^ PBMC, but were still well above the control mice, 30 spots/10^6^ PBMC respectively (Figure 3). The data demonstrates that AAV-immunized mice harbor spike-specific T-cells and when reposed to antigen produce interferon-γ.

**Figure 3.**
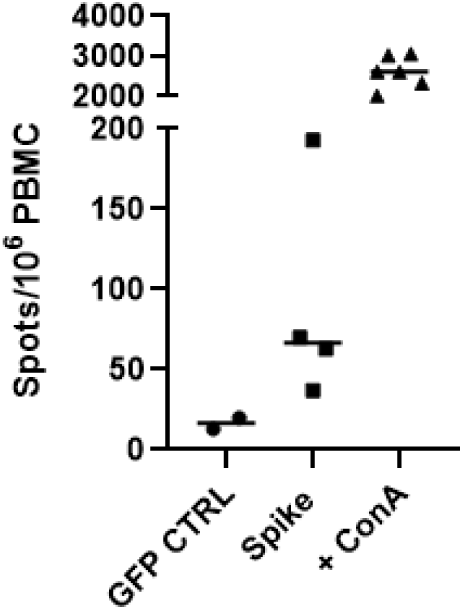
Levels of IFN-γ in Splenocytes Obtained from Mice Transduced with AAV5-GFP or AAV5-spike and Challenged with a Mixture of Spike peptides. ELISpot assay was used with splenocytes extracted from animals 16 weeks after vaccination and stimulated with SARS-CoV-2 spike peptides. The levels of IFN-γ positive PBMCs from the AAV-GFP (GFP CTRL) and AAV5-Spike (Spike) transduced mice. As shown on the left side of the figure, splenocytes were treated with ConA (+ ConA) as positive control for T-cell activation. The horizontal bars represent the median value in each group.

Memory T-cells were also evaluated by gated FACS analysis by staining for several antibody markers including CD4, CD8, and CD44 bright. However, the analysis at the 4-month time point did not reveal any difference in T-cell populations derived from the AAV spike transduced mice compared to the AAV5-GFP mice (data not shown). Repeat immunization did not result in a detectable expansion of the memory T-cell population (Supplemental Figure 3).

### AAV5 DNA Persistence at the Site of Administration

AAVs have a record of persistence of transduction implying the presence of active transgene after vector administration. In case of vaccine gene therapy, presence, or absence of transgene DNA in the site of administration can reveal whether the antigen is continuously expressed. Therefore, the reaction of the immune system may have relevant outcomes. A separate group of mice was used to examine the IM AAV administration tissue site for the presence of AAV vector DNA. Muscle tissue from the IM site of administration was collected, DNA extracted and quantified by qPCR. As shown after one week (Figure 4), a high level of AAV DNA was present. However, a dramatic drop was observed by week four and a further decrease was observed at week 8-12 showing approximately only 100 viral particles per ug cellular DNA.

**Figure 4.**
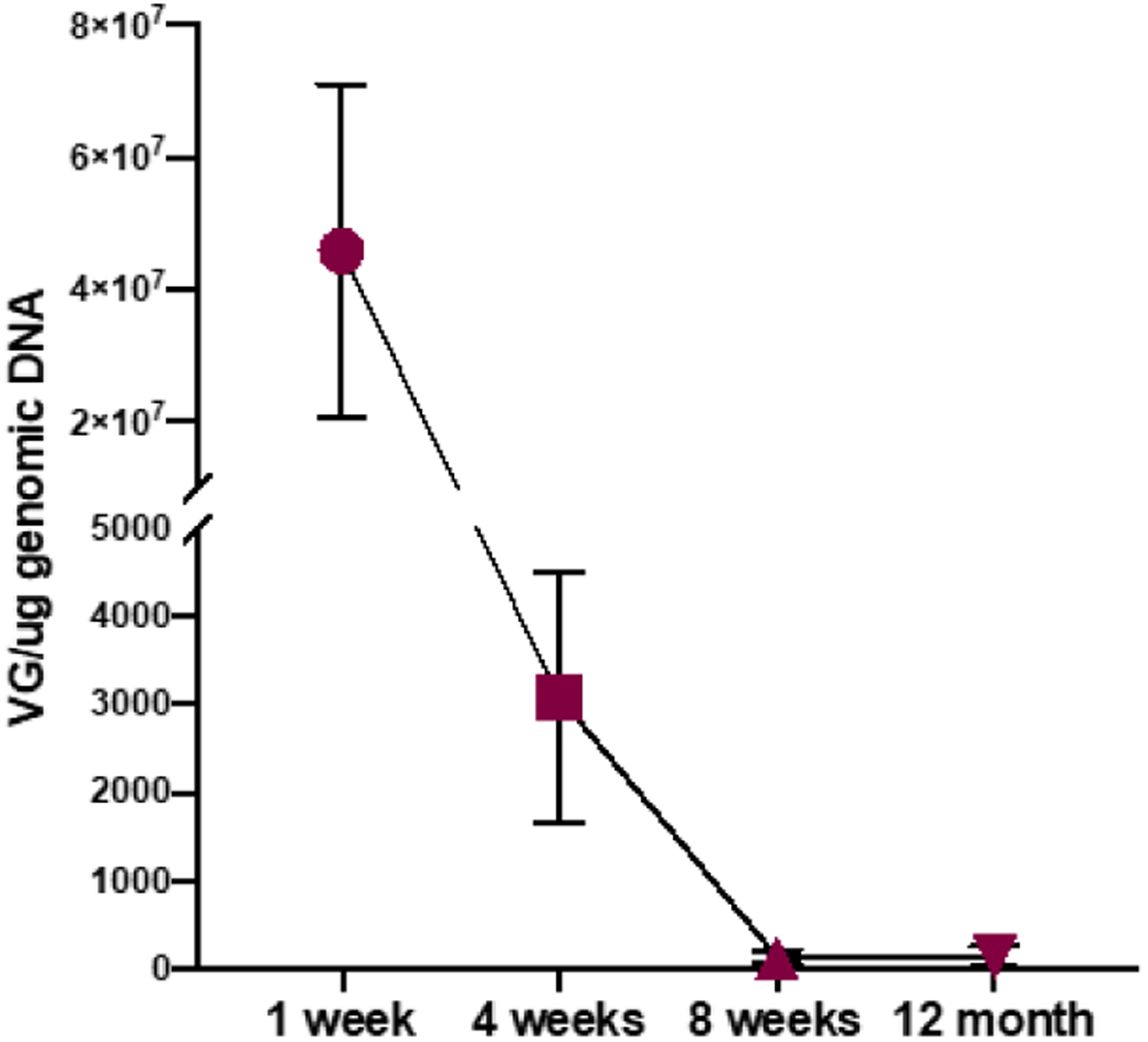
Longitudinal Analysis of the Amount of Detectable AAV5-Spike DNA at the Site of IM inject. Mice administered IM AAV5-Spike were monitored over time to determine the turnover of the AAV5 at the site of muscle injection. Biopsied material was then subjected to qPCR as described in the material and methods.

## Discussion

Our study adds to the existing literature establishing the AAV system as a potent technology for generating vaccine-mediated immunity against SARS-CoV-2 and likely other respiratory viruses. We show that three AAV serotypes, AAV5, AAV44.9, and AAV12 were highly effective in generating long-lasting vaccine-associated humoral immunity against the SARS-CoV2 spike protein. At the initiation of this work, little was known about which SARS-CoV-2 spike protein would be the most effective immunogen. We generated the full-length spike and spike trimer stabilized to match antigens used for human vaccination [18] and unsurprisingly found that the subtle amino acid differences between the full-length and spike-stabilized mutant trimer forms generated similar antibody levels and neutralizing activity in mice. Other studies using AAV technology have successfully employed smaller fragments of the SARS-CoV-2 spike protein, focusing on the receptor-binding domain, to effectively vaccinate and generate both humoral and cellular immunity [10, 12–14, 17]. In one study, AAV delivery of the full-length spike protein compared to an AAV-RBD domain only protein demonstrated markedly lower antibody levels after day 100 [10], highlighting the importance of the design of the vaccine antigen and that the full-length spike need not be used to elicit an effective vaccine.

Two AAV spike vaccine studies mainly focused on antibody production during the first several months after administration [14, 22]. Our work extends these observations and demonstrates that spike antibodies remain high for over a year with both IM and IN administration. Unlike other AAV studies which typically looked at two weeks post-administration, we also document long-term cellular immunity at 16 weeks with ELISpot T-cell memory production of interferon-γ after spike peptide stimulation. In contrast to all the other studies that detected immune subset changes at 2 weeks post AAV administration, the changes in immune subsets at the 16-week time point were not detected in our samples. Additionally, we provided evidence for the lack of amplifiable AAV-Spike DNA in the tissue after one week, yet we observed long-lasting immunity from a single dose of AAV administration. The observed high levels of spike antibody observed in AAV-transduced mice are likely maintained by memory B-cells or long-lived plasma B cells. These results contrast with other SARS-CoV-2 spike vaccine technologies such as adenovirus expression and mRNA-vaccine delivery which are much more transient and therefore require boosting. On the other hand, one potential disadvantage of AAV technology is the robust and sustained antibody expression, which may lead to desensitization and downregulation of the immune response. Nevertheless, the long-lasting vaccine immunity warrants further investigation and highlights the possibility of using the AAV system for long-lasting vaccines for infectious diseases and even cancer.

IM administration is generally not thought to offer an effective route to provide mucosal immunity against SARS-CoV-2 for the respiratory tract using adenovirus and mRNA vaccine delivery technologies. However, it is possible that the high level of immunity generated by AAV spike expression does provide protection against infection as documented in a macaque rhesus challenge studies [11]. Our study confirms the efficacy of IN administration for SARS-CoV-2 spike vaccination but noted no significant difference in antibody levels or neutralization activity with IM administration. More recently, inhalation delivery has been shown to be even better than IN administration for inducing mucosal immunity against viruses [23, 24]. The likely reason is that more of the vector is targeted to the lung as shown by a mouse study, which found that IN administration of labeled nanoparticles resulted in only 28% reaching the lungs and most reaching the stomach [25]. In contrast, inhalation administration of nanoparticles resulted in 85-95% reaching the lungs. It would be interesting to test whether even lower doses of AAV-spike delivered by inhalation could be an effective strategy for sustained mucosal immunity. In conclusion, our results along with the other published studies support the finding that the AAV, independent of the AAV used or the route of administration, produces a highly robust humoral and cellular immunity against the SARS-CoV-2 spike protein.

## Methods

### Design of rAAV Vectors for SARS-CoV-2 Spike Protein Expression

Two different AAV serotypes, AAV5 and AAV44.9, were chosen as vectors to express the recombinant SARS-CoV-2 spike antigen to induce vaccine immunity (Supplemental Figure 1). Two different recombinant SARS-CoV-2 spike proteins, the entire ectodomain (amino acids 1-1208) of the spike protein, as well as a Spike-trimer stabilized protein (to lock in pre-fusion conformation to allow for optimal RBD antigenicity), were utilized based on published the published sequence [18]. Recombinant AAVs were generated in 293T cells by calcium phosphate transfection with four plasmids, which included the pAd12 adenovirus helper plasmid containing VA RNA and coding the E2 and E4 proteins; AAV helper plasmids containing the AAV rep and the AAV5 or AAV44.9 capsid genes respectively; and vector plasmids including the AAV2 inverted terminal repeats flanking the two different spike gene expression cassettes. The mixture was transfected into five 15-cm-diameter plates and contained 32 μg of each plasmid DNA except for 32 mg DNA except for using 72 μg of the pAd12 plasmid. Forty-eight hours after transfection, the plates were harvested, and a crude viral lysate was generated by one freeze-thaw cycle. The lysate was treated with 0.5% deoxycholic acid along with 100 U/mL DNase (Benzonase, Sigma) for 30 min at 37°C. Low-speed centrifugation was used to collect vector particles present in the clarified lysate, which were further purified by cesium chloride gradient ultracentrifugation. Titers of the AAV were determined by quantitative real-time PCR (Applied Biosystems, Foster City, CA, USA). The vector doses were dialyzed against PBS using a 2ml Float-A-Lyzer G2 1000 kD dialysis device (Spectrum Labs). Vector was concentrated using a centrifugal filter unit (Amicon Ultra 0.5mL 100k). AAVr particles were quantified by qPCR. One microliter of diluted recombinant preparation was added to a PCR containing 1×SYBR Green Master Mix (ABI) and 0.25 pmol/μL forward and reverse primers. Amplification was detected using an ABI 7700 sequence detector (ABI). Specific primers CMV forward 5′-CATCTACGTATTAGTCATCGCTATTACCAT-3′, CMV reverse 5′-TGGAAATCCCCGTGAGTCA-3′. Following denaturation at 96°C for 10 min, cycling conditions were 96°C for 15 s, 60°C for 1 min for 40 cycles. The viral DNA in each sample was quantified by comparing the fluorescence profiles with a set of DNA standards to determine the number of DNAase-resisting particles (DRP).

### Animal Studies and *In Vivo* AAV Intramuscular and Intranasal Transduction

All mouse animal studies were approved by the Animal Care and Use Committees of the National Institute of Dental and Craniofacial Research. Female BALB/c mice were obtained from Charles River Laboratories. Several series of mouse transduction experiments in which AAV5 was administered intramuscularly were performed to collect the needed serum and other tissues. In one series of mouse experiments, groups (*N* = 4-6) of 8-week-old female BALB/c were intramuscularly administered AAV5 spike and AAV5-spike trimer at 5.10^10^ DRPs. Serum samples from the mice were then obtained at 2, 4, 8, and 16 weeks after transduction, as well as additional time points post-transduction were examined for spike antibody levels and capacity for neutralization. Another set of AAV-5 spike mice was sacrificed at 4 months post-transduction to look at cellular immunity by ELIspot and FACS. Lastly, a parallel set of mice who received AAV5 spike intramuscularly were sacrificed at 1, 4, 8 weeks and 12 months later and their muscle tissue were examined for the presence of AAV5 genomic DNA by quantitative PCR.

For the AAV nasal administration experiments, a group (*N* = 5) of 8-week-old female BALB/c mice was administered intranasally with 5 ×10^10^ DRP in 150 µl of PBS of AAV5-spike, AAV5-spike-trimer, AAV44.9-Spike or AAV12-spike. Serum samples were obtained over time by retro-orbital bleeding collection. Sera was used to measure anti-spike antibody levels and testing for spike neutralization.

### Determination of SARS-CoV-2 Antibody levels

The luciferase immunoprecipitation systems (LIPS) immunoassays were used to assess antibody levels against SARS-CoV-2 spike protein as described [19, 20]. In these and other studies, a SARS-CoV-2 spike protein fragment (aa 1-538) was used to measure levels of antibody. Batched serum samples collected over the different time points were tested in real-time for spike antibody levels. LIPS immunoassay testing on mice sera was standardized using seropositive human sera from natural SARS-CoV-2 infection and from sera of COVID-19 vaccine immunized individuals.

### Detection of Neutralizing SARS-CoV-2 Antibody

To assess neutralizing antibodies, the cPass SARS-CoV-2 Neutralization Antibody Detection kit (GeneScript, Piscataway, NJ) was used according to the manufacturer’s instructions. Briefly, samples, along with positive and negative controls, were diluted at a ratio of 1:9 with sample dilution buffer [21]. Briefly, precoated recombinant hACE2 protein attached to a microtiter plate is incubated with antisera and HRP-labeled RBD (HRP-RBD). When neutralizing antibodies against SARS-CoV-2 spike protein are present in sera, block of specific protein-protein interaction between ACE2 and HRP-RBD occurs, hence reducing the chromogenic reaction. To initiate the assay, serum samples were diluted 1:10 and preincubated with HRP-labeled RBD for 15 min at 37° C. The mixture is then added to microtiter plates pre-coated with the human ACE2 protein. After several washing steps with PBS, the signal was measured by adding TMB solution and then with stop solution using a microtiter plate reader (Molecular Devices SpectraMaxi3, Molecular Devices, San Jose, CA) to read absorbance. Sera samples with more neutralizing antibodies showed lower signal intensities, in which the inhibition percentage was calculated using the equation: (1-(OD value of sample / OD value of negative control)) × 100%.

### Cellular Immunity Measuring IFN**γ** Production and Memory T-Cells

Cellular immunity was assessed in mice following five months post-transduction. For these studies, 300K splenocytes were isolated from each of the two AAV5-GFP or four AAV5-spike-treated mice and were plated into IFNγ coated ELIspot wells. Cells were stimulated in the following conditions for 18-24 hours: cells alone, 1 μg/mL SARS-CoV-2 Pepmix, or 2 μg/mL ConA. Wells containing cells were washed with PBS before a biotinylated detection antibody was added for two hours at room temperature. Next PBS was used to wash the wells and Streptavidin-ALP was added and incubated for 1 hour at room temperature. Further PBS wash followed, then spot development was initiated by adding BCIP substrate. Lastly, wells were washed with water and dried. Spots were quantified by counting on an Immunospot S6 universal analyzer.

The relative number of memory T-cells was measured by the FACS gating strategy. Briefly, 500K lymphocytes were obtained from each of two AAV5-GFP or four AAV5-spike transduced mice. Following washing with PBS, the cells were incubated with the following antibodies: AF700 CD8, PC5 CD4, BV650 CD44, and APC CD62L. Staining for CD44 bright and either CD4 or CD8 was used to detect memory T-cells.

### Determination of Residual AAV in Tissue

To determine the persistence of the AAV5-spike DNA in IM-treated mice, a parallel set of AAV5-spike-treated mice were biopsied at the injection site. DNA tissue extraction was performed using a Promega Genomic DNA extraction kit according to the manufacturer protocol and quantified by qPCR as described above for AAV recombinant quantification. AAV copy number DNA per cell was established by calculating the ratio ρgs cellular template divided by the viral copy number obtained from the qPCR reaction respectively, assuming that x ρgs DNA equals 100 total cell DNA. Untreated tissue DNA was used as a control. The limit of detection (LOD) by qPCR was 10 particles.

## Data availability

Primary data associated with figures and tables are available from the corresponding author on request.

## Acknowledgments

The authors would like to thank Vincent Eun and Yunju Jung for comments and suggestions. This work was supported by the Division of Intramural Research, NIDCR/NIH (1ZIADE000695, JAC).

## Author Contributions

Y. Ji, G. Di Pasquale G., H. Yin P.D. Burbelo and J.A. Chiorini contributed to the manuscript conception and design. Y. Ji, G. Di Pasquale, C. Zheng, S. Afione, T. Esperanza and P.D. Burbelo performed data generation, analysis and interpretation. Y. Ji, G. Di Pasquale, P.D. Burbelo and J.A. Chiorini drafted and critically revised the manuscript. All authors gave final approval and agreed to be accountable for all aspects of the work.

