## Supplemental figures for "Intranasal AAV Vaccination of SARS-CoV-2 Induce Strong and Sustained Neutralizing Antibodies in Mice"

**Supplemental Information**

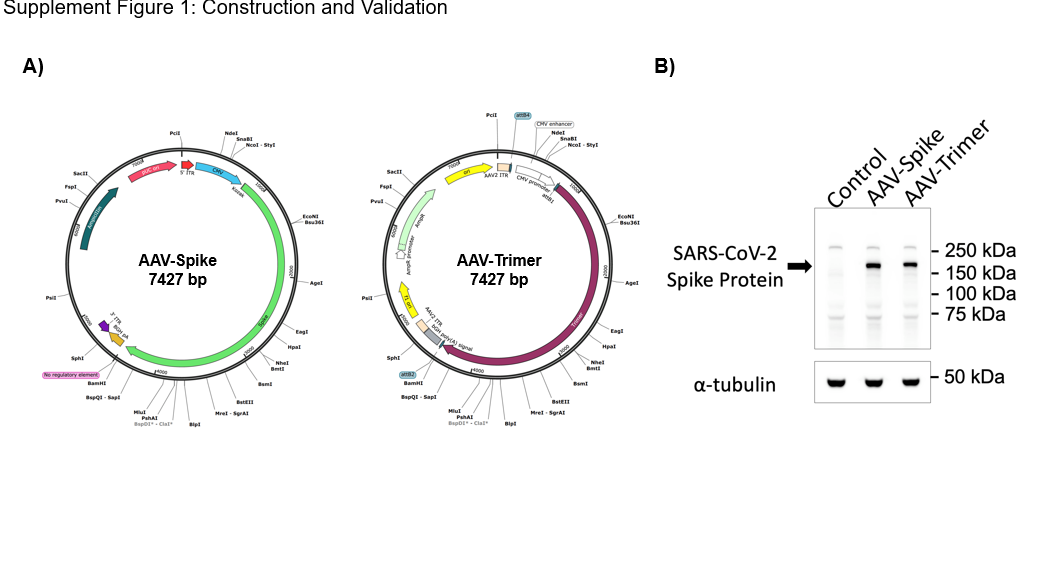

**Supplemental Figure 1. Design and in vitro characterization of AAV vectors for SARS-CoV-2 spike and Spike-stabilized trimer form.** **A)**. Two different recombinant SARS-CoV-2 spike proteins, the entire ectodomain (amino acids 1-1208) of the spike protein, as well as a Spike-trimer stabilized protein (to lock in pre-fusion conformation to allow for optimal RBD antigenicity) were utilized. **B).** Expression Spike protein after transfection. Cell lysate was harvested and Western blotting was performed with a rabbit anti- SARS-CoV-2 spike commercial antibody (Invitrogen).

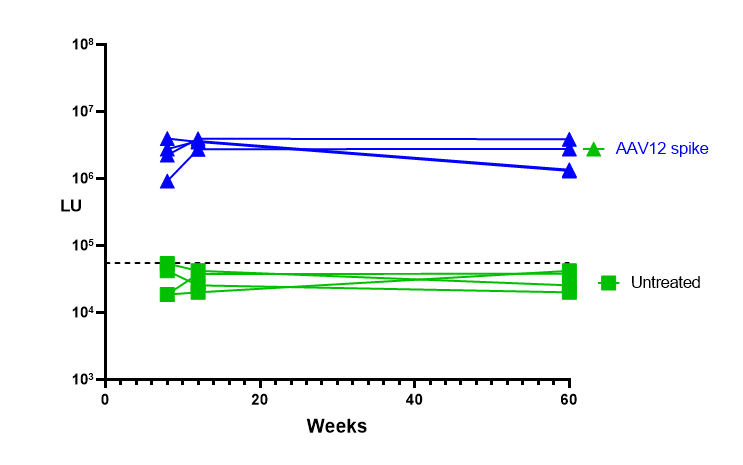

**Supplemental Figure 2. SARS-CoV-2 Spike Antibody Levels by LIPS in AAV12-spike mice by IN Administration.** Each line represents a value from each serum from mice over time. As control, untreated serum mice were also quantified and shown individually. Antibody levels are expressed in relative light units (LU) on a log scale. Dashed line separate value of treated and untreated mice.

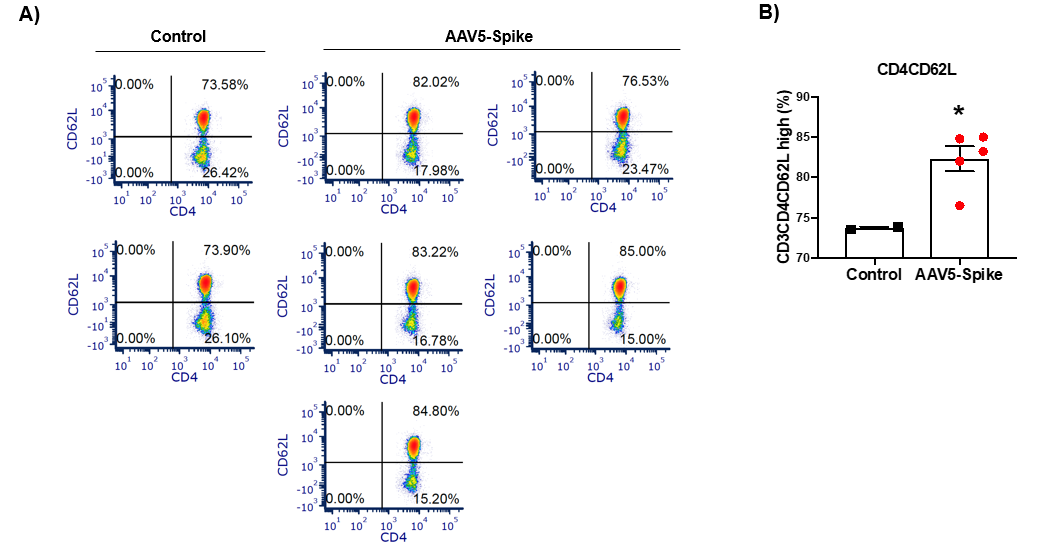

**Supplemental Figure 3. Determination of Memory T cells in AAV-spike Administered Mice.** As described in the material and methods, FACS analysis was used with a panel of immune surface antibody markers and analyzed by FACS to assess whether changes in immune cells population changes after AAV-spike immunization. No significant changes in cell populations were observed in the mice receiving AAV-spike.
